# Mapping global transcript isoform landscapes of influenza A virus infection and interferon stimulation reveals RNA-binding protein networks associated with immune remodeling

**DOI:** 10.64898/2026.09.22.753599

**Authors:** Simon Boudreault, Rosa Hernansaiz-Ballesteros, Attila Gábor, Gurdeep Singh, Steven F. Baker

**Affiliations:** Children’s Hospital of Eastern Ontario Research Institute, Ottawa, ON, Canada; Department of Biochemistry, Microbiology and Immunology, University of Ottawa, Ottawa, ON, Canada; Tempus AI, Inc., Chicago, IL, USA; Department of Ophthalmology and Visual Sciences, University of New Mexico School of Medicine, Albuquerque, NM 87131, USA; Department of Cell Biology and Physiology, University of New Mexico School of Medicine, Albuquerque, NM 87131, USA; UNM Comprehensive Cancer Center, Albuquerque, NM 87131, USA; Department of Systems Biology, George Mason University, Manassas VA 20110, USA; Infectious Disease Program, Lovelace Biomedical Research Institute, Albuquerque, NM, USA; Department of Molecular Genetics & Microbiology, University of New Mexico School of Medicine, Albuquerque, NM, USA; Department of Microbiology and Immunology, Loyola University Chicago, Maywood, IL, USA

## Abstract

Infected cells rapidly adapt their transcriptional profile to upregulate genes that limit pathogen replication and promote inflammation. Viruses also induce widespread changes in host gene expression, and both host and virus further remodel the transcriptome through targeted mRNA degradation, modulation of alternative splicing, and alterations at transcript 5’ and 3’ ends. The complexity of the response to infection makes it difficult to understand which transcriptional changes are driven by the virus and which reflect the host antiviral program. Short-read RNA sequencing has been the primary tool for studying these changes, but its inability to resolve fulllength transcript isoforms limits isoform switching-based analyses, narrowing our understanding of how transcript usage shapes immunity. Here we profiled the host transcriptome in response to interferon-β treatment or influenza A virus infection using both short- and long-read RNA sequencing analyzed with the SQANTI3 pipeline. Because isoform switching and transcript abundance change are distinct regulatory events, we focused our analysis on differential transcript usage to uncover a layer of transcriptome remodeling that standard expression analysis cannot detect. Pathway analysis revealed distinct regulons for genes undergoing transcriptional upregulation compared with those undergoing isoform switching. These data provide a rich resource for querying the isoform changes underlying shared and distinct transcriptional programs triggered by viral infection or innate immune stimulation. To identify putative mechanisms coordinating these stimulation-dependent isoform changes, we integrated 232 eCLIP datasets and found a small set of RNA-binding proteins localized at these isoform switches, including DDX3X. Comparing the responses to interferon and to virus highlighted RNA-binding proteins associated with infection-dependent immune remodeling. Altogether, these results demonstrate the value of hybrid sequencing for deciphering the post-transcriptional regulatory code active during the antiviral response.

## INTRODUCTION

Infected cells deploy a variety of programs to fight viral infection. The first line of cellular defense is a two-pronged system consisting of innate immune sensing and the subsequent interferon response. Upon detection of pathogen-associated molecular patterns produced during viral replication, pattern recognition receptors such as RIG-I, MDA-5, and cGAS trigger a signaling cascade that culminates in IRF3 phosphorylation and nuclear translocation, induction of type-I interferon, notably IFN-β, and its secretion into the extracellular milieu (1). Secreted type-I interferons can act in an autocrine or paracrine fashion, stimulating both the infected cell and bystander cells through the IFNAR receptor to drive expression of interferon-stimulated genes (ISG). These ISG are responsible for fighting viral infection and protecting bystander cells from further infection (2,3). Since most viruses trigger the interferon response to some extent, separating the direct effects of the virus from those of the host immune response remains challenging (4).

Viruses induce widespread transcriptomic changes during infection, including disruption of transcription termination (5), alterations in polyadenylation (6), and modulations in alternative splicing (7–10). For Influenza A virus (FLUAV hereafter), widespread changes in alternative splicing have been previously described (11,12) and linked to several RNA-binding proteins (RBPs), such as hnRNP K, NS1BP, RED, and Smu1 (11–13). However, FLUAV has a more profound and wide-ranging impact on the host cell transcriptome. For example, the NS1 protein inhibits 3’ end processing by interacting with CPSF4, preventing cleavage and polyadenylation of cellular mRNAs, which are then retained in the nucleus (14–16). FLUAV also triggers targeted decay of host transcripts through the PA-X protein, a virus-encoded endoribonuclease (17–19). Finally, FLUAV triggers transcriptional readthrough through defective Pol II termination linked to the cellular stress response (20). Most of these transcriptomic alterations shape the outcome of viral infection, underscoring the crucial roles they play in FLUAV-host interactions (12,15,16,19,21).

RNA-binding proteins (RBPs) play a key role in the life cycle of mRNAs. They control transcript fate by guiding splicing decisions, dictating RNA localization, accessibility, and interactions, and shaping mRNA half-life (22). In the context of innate immunity, the roles of some RBPs are well understood, notably the control of alternative polyadenylation by core polyadenylation machinery proteins (23). RNA stability and decay also contribute to finetuning the innate immune response through regulatory feedback loops involving RBPs such as TTP, ZFP36L1, and ZFP36L2 (23–27). However, knowledge about how RBPs control alternative splicing and isoform switching during innate immune sensing remains sparse (23). A gene can toggle its predominant isoform without changing its total RNA output. This makes isoform switching invisible to standard expression analysis, leaving a major gap in understanding the transcriptomic response to infection. Previous studies have implicated SRSF7 in inducing IRF7 expression at the transcriptional level, and hnRNP M controls the splicing of some cytokines (28–30). In the case of viral infection, multiple RBPs are usurped by viruses, usually to benefit replication (31–36). For example, sponging of HuR by the 3’-UTR of Sindbis virus alters mRNA stability, splicing, and polyadenylation of host transcripts (37). Nonetheless, for most of these RBPs, the impact of their co-option by viruses on host-cell splicing remains incompletely understood, and the specific RBPs that orchestrate host immune remodeling during viral infection remain poorly defined.

The advent of short-read RNA sequencing sparked a transcriptomic revolution, allowing researchers to broadly probe the RNA landscape of cells and tissues (38). However, short-read sequencing has intrinsic weaknesses, most notably read length, which precludes full-transcript resolution. Long-read sequencing was developed to address these limitations, but these technologies still lack the sequencing depth needed to cover low-abundance transcripts, and some suffer from high error rates (39). In most cases, full transcriptome resolution remains ill-defined under non-homeostatic conditions. SQANTI3 is a bioinformatic pipeline that addresses the weaknesses of both sequencing modalities and generates high-resolution full transcriptomes (40). By integrating long- and short-read RNA-seq, SQANTI3 enables precise curation, definition, and quantification of expressed transcripts. This method has already been successfully applied to cancer, reproduction, immunity, and plant biology (41–44). The integration of bioinformatic and data-driven approaches that leverage publicly available datasets has proven central to advancing our understanding of fundamental regulatory mechanisms (45–47). For example, pathway analysis can identify regulons that drive biology (48,49), and the ENCODE consortium provides a rich source of data encompassing KD-RNA-Seq, eCLIP, and Bind-N-Seq to investigate RBP-mediated regulation (22).

In this study, we use long- and short-read sequencing during FLUAV infection or IFN-β stimulation to resolve the shared and distinct transcriptomic changes underlying each response. Using SQANTI3, we profile stimulus-induced isoform changes and identify oppositely regulated transcripts and genes with differential transcript usage. We then use pathway analysis to define the regulons associated with each condition, and integrate ENCODE eCLIP data to nominate RBPs, such as DDX3X, positioned at the switching events that are both shared with and distinguish FLUAV infection from IFN-β stimulation. This resource maps the isoform landscape of innate immune stimulation and viral infection and generates mechanistic hypotheses about the RBPs that shape it. Functional validation of individual RBP-isoform pairs is the subject of ongoing work.

## RESULTS

### A hybrid long- and short-read transcriptome resolves the isoform landscape of infected and interferon-stimulated lung cells

Full transcriptome resolution remains ill-defined in numerous stimulation conditions, notably during viral infection and the interferon response. To address this gap, we leveraged our short-read (21) and long-read RNA sequencing data (50) together with SQANTI3 to generate a high-resolution transcript map of human A549 lung cells upon influenza A virus (FLUAV) infection or IFN-β stimulation (**Figure 1A**). Upon curation and classification of the collapsed isoforms, potential artifacts were removed and the rescue module was applied in automatic mode to avoid discarding genuine transcripts that fail orthogonal validation (**Figure S1A**). A complete description of the analysis pipeline is provided in the Methods. The resulting hybrid transcriptome comprised 10,185 genes and 34,000 transcripts, of which 11,713 were identified as artifacts leaving 22,287 bona fide full length transcripts (**Figure S1B**). SQANTI3 annotates isoforms by comparing their splice junctions to the reference transcript (**Figure S1B**). If all splice junctions are perfectly matched to the reference, the isoform is categorized as a full splice match (FSM). This is the only category of known isoforms. All isoforms not categorized as FSM are novel. If internal splice junctions are matched to the reference but some 5’ or 3’ junctions are absent, the transcript is classified as an incomplete splice match (ISM). If the transcript contains a new combination of known splice sites, it is classified as novel in catalog (NIC). If the isoform contains at least one novel donor or acceptor site, it is classified as novel not in catalog (NNC). Identified isoforms were primarily full-splice-match (FSM), followed by incomplete-splice-match (ISM) and novel-in-catalog (NIC). The remaining categories were the least populated after artifact removal (**Figure S1C**). The median transcript length was 2.5 kb (**Figure S1D**). For FSM isoforms, the transcription start site (TSS) was generally close to the annotated TSS (**Figure S1E**). For ISM isoforms, the TSS was usually more than 5 kb from the annotated TSS (**Figure S1F**).

**Figure 1.**
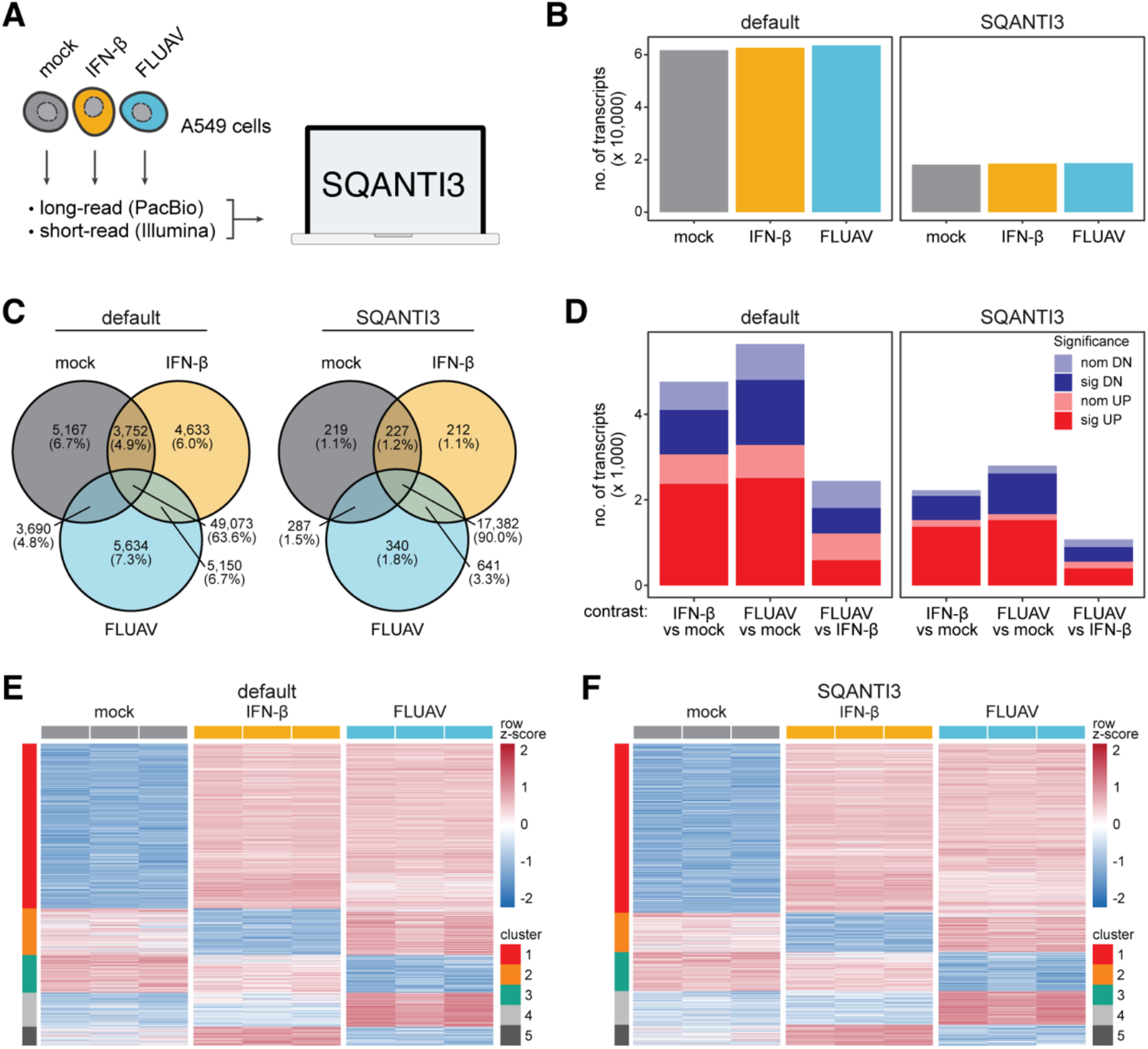
SQANTI3-powered high-resolution analysis of transcript expression in stimulated human lung cells. **A**, Experimental design. RNA libraries were prepared from human lung A549 cells mock or type-I interferon (IFN-β)-treated, or influenza A virus (FLUAV)-infected for long- and short-read RNA-sequencing. SQANTI3 combines these sequencing data to create a full-length transcriptome against which short reads were then aligned for downstream quantitation. **B**-**F**, Comparison of short-read RNA-sequencing data aligned to hg38 and quantified against Ensembl (default) or SQANTI3-informed transcript models. **B**, Number of unique transcript isoforms detected. **C**, Overlap of transcript isoforms across conditions. **D**, Number of significant (sig, *P*-adj < 0.01) and nominally (nom, *P* < 0.01) up- and down (DN)-regulated transcripts. **E**-**F**, Differentially expressed transcripts were clustered across biological replicates and between conditions through k-means clustering.

We next compared differential transcript expression when aligning matched short-read RNA-seq data to hg38 using standard Ensembl-defined transcript models versus our full-length A549 transcript assembly. Aligning to the SQANTI3 transcriptome yields roughly one-third as many transcripts as standard processing, suggesting that SQANTI3 maps short reads more stringently to expressed isoforms (**Figure 1B**). Only 63.6% of the transcripts detected using short-read data alone are shared across the three conditions, suggesting substantial variability in short-read-derived isoform identification even within the same cell line. This shared fraction increases to 90% with SQANTI3, underscoring its high signal-to-noise ratio for detecting bona fide transcripts and removing artifacts (**Figure 1C**). The proportion of transcripts unique to one condition or shared across only two conditions does not exceed 3.3%, indicating that most bona fide isoforms are reproducibly detected across all three conditions. However, the dramatically high number of such unique condition-specific isoforms identified with short-read data alone suggests that short-read based computational transcript reconstruction generates isoform artifacts. Differential transcript expression (DTE) analysis using DESeq2 (51) showed that SQANTI-defined annotations yielded a higher proportion of differentially expressed transcripts compared to the default annotation (**Figure 1D, Table S1A**). Selecting the top 500 differentially expressed transcripts using standard (**Figure 1E**) or SQANTI-defined annotations (**Figure 1F**) allows clear clustering of transcripts by treatment group, suggesting that both methods capture the transcript signature unique to viral infection or IFN-β stimulation. As previously established, SQANTI3-defined annotations outperform default annotations in generating cell type- and state-specific high-confidence transcript models. SQANTI3-defined annotations also recovered typical infection-specific differentially expressed transcripts (**Figures S2A-C**). These results confirm the generation of a high-resolution transcript map of A549 cells upon FLUAV infection and IFN-β stimulation.

### Transcript abundance reveals coherent interferon programs alongside discordantly regulated pathways

We next parsed DTE results to identify pathways regulated during infection and the interferon response. Because a pathway can be remodeled in two distinct ways, over-represented KEGG and Reactome terms were ranked by two complementary statistics. We used a signed test, which sums the log_2_ fold change of member transcripts, to identify pathways whose transcripts move coherently in one direction. We also used an absolute test, which sums the magnitude of transcript movement irrespective of sign, that identifies ‘discordant’ pathways carrying substantial transcript movement whether that movement is directionally coherent. This analysis revealed that the directionally coherent pathways recovered the expected biology. For example, upon IFN-β treatment, *Interferon Signaling, Interferon alpha/beta signaling, Interferon gamma signaling, and Signaling by Interleukins* were upregulated, together with other terms linked to viral infection and immune signaling, as expected given the broad transcriptional response induced by IFNAR signaling (**Figure 2A**). Similarly, the FLUAV infection produced the same coherently upregulated interferon terms, consistent with robust activation of the interferon response by the virus, along with coherent downregulation of respiration (*Respiratory electron transport, Aerobic respiration and respiratory electron transport*) and translation (*Eukaryotic Translation Elongation, Ribosome, Eukaryotic Translation Initiation*) (**Figure 2B**). Direct comparison of FLUAV to IFN-β isolated these virus-specific effects from the interferon program. Translation terms (the same as FLUAV vs mock), respiration, and reactive oxygen species pathways (*Oxidative phosphorylation, Chemical carcinogenesis – reactive oxygen species, Metabolism of amino acids and derivatives*) remained downregulated, with no interferon terms recovered, as expected (**Figure 2C**).

**Figure 2.**
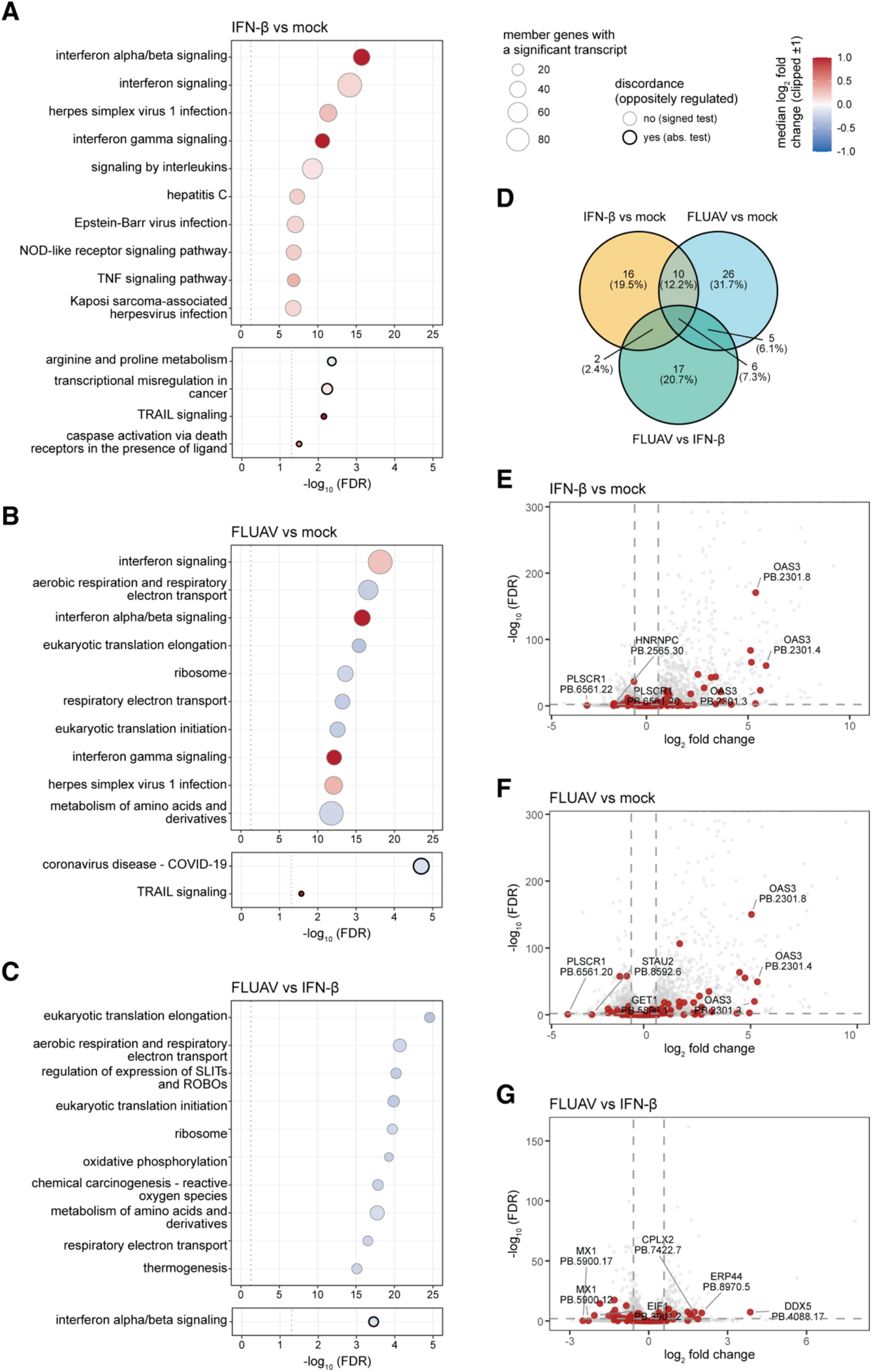
Pathways and genes with discordantly regulated transcripts. **A**-**C**, Pathways of genes that are enriched to contain differentially expressed transcripts. Directionality (log_2_ fold change, log2FC) determined by the sum of transcript movement within a given pathway. Pathways ranked by a signed test to identify directional regulation or an absolute (abs) test to detect magnitude of change regardless of direction. **D**, Genes that encode oppositely regulated transcripts between indicated conditions. **E**-**G**, Differentially expressed transcripts between indicated conditions. Red circles indicate oppositely regulated transcripts from **D**. Top three differentially expressed transcripts indicated with their PacBio isoform ID number.

Since both FLUAV and IFN-β stimulates the interferon response, we wanted to clarify their respective induction of this pathway. To ascertain the respective impact of FLUAV and IFN-β on the interferon response, we measured the induction of IFN-β (*IFNB1*) and interferon-stimulated genes (*IFIT1, IFIT2, EIF2AK2*) using qPCR (**Figure S2D**). As expected, IFN-β treatment only has a minimal impact on induction of its own gene under these experimental conditions, whereas FLUAV robustly triggers a 25-fold induction in *IFNB1*. With the exception of *IFIT2* that is similarly induced by FLUAV and IFN-β, *IFIT1* and *EIF2AK2* are induced to a higher extent with IFN-β than with FLUAV, in agreement with our understanding of FLUAV antagonism of the interferon response (53). These results confirm that, in agreement with our GO analysis, FLUAV triggers a robust IFN response, albeit to a less extent than direct IFN- β stimulation.

Discordant pathways contained terms linked to programmed cell death (*TRAIL signaling, Caspase activation via Death Receptors in the presence of ligand*), ranked high under the absolute test but did not reach significance under the signed test (**Figure 2A-C**). These pathways were therefore not simply switched on or off during infection. Transcripts within them were being raised and lowered simultaneously, an abundance-level signature expected if the underlying regulatory event were a change in which isoform a gene produces rather than a change in how much RNA that gene makes. This is consistent with the established importance of isoform pairs encoding proteins of opposing function in cell death regulation (54).

Discordance at the pathway level implies discordance at the gene level, so under the same contrasts we searched for genes encoding oppositely regulated transcripts (ORTs), defined as genes for which at least one transcript is significantly regulated in one direction while at least one other transcript is significantly regulated in the opposite direction. Across the three contrasts, 82 unique genes met this definition, accounting for 111 gene-contrast instances (mock vs IFN-β, 34; mock vs FLUAV, 47; FLUAV vs IFN-β, 30; **Figure 2D, Table S1B**). Of these, 23 genes were recovered in at least two contrasts and 6 in all three. Whereas overall DTE was strongly biased toward overexpression, ORTs were distributed across the full range of fold changes observed, including transcripts with modest individual effect sizes (**Figure 2E-G**). ORTs therefore provide direct, gene-level confirmation of the discordance inferred from the absolute pathway test. ORT detection is nonetheless constrained by the abundance framework that defines it. A gene qualifies as an ORT only when two or more of its transcripts each independently clear a significance and fold-change threshold in absolute abundance. Genes that redistribute output between isoforms without either isoform crossing that threshold, and genes whose total output is unchanged, are excluded by construction. ORTs are thus the subset of discordant regulation that happens to be visible in abundance space, not its full extent.

Finally, we asked whether the pathways defined above via DTE could be used to nominate the RBPs responsible for these changes based on their representation within the enriched pathway. Ranking the 1,371 retained RBPs (pared down from (55), see methods) by their representation in the DTE-derived pathway universe (557 genes with at least one pathway membership) yielded a compact RBP shortlist primarily composed of factors associated with nucleic acid sensors and downstream signaling proteins such as TLR3, MDA5, PKR, and eIF2α (**Figure S2E**). mRNA maturation factors were absent from the list. Because the pathway universe was built from DTE, which measures net transcript abundance, the ranking preferentially recovers regulators of transcriptional output and is structurally blind to the machinery that redistributes output between isoforms. Taken together, DTE established the coherent transcriptional programs driven by FLUAV and IFN-β, demonstrated through discordant pathways and ORTs for which a substantial component of the response is not directional, and showed that abundance space alone can neither resolve that component nor identify the proteins controlling it.

### Differential transcript usage resolves isoform switching that abundance analysis cannot detect

A gene can shift which isoform it predominantly produces without changing its total RNA output. This switch can carry profound functional consequences because different isoforms may encode proteins with distinct or opposing activities. Standard abundance analysis is blind to this layer of regulation, making differential transcript usage (DTU) the appropriate tool for detecting how infection truly remodels the transcriptome. DTU analysis was thus performed using IsoformSwitchAnalyzeR (see methods). FLUAV infection and IFN-β treatment each drove widespread isoform switching as determined by the change in isoform fraction (ΔIF) in A549 cells: 377 genes across contrasts, with 160, 207, and 145 significantly switching genes for mock vs IFN-β, mock vs FLUAV, and FLUAV vs IFN-β, respectively (**Figure 3A-C, Table S1C**). DTU analysis also recovered more than half of the ORTs identified by DTE, while capturing many additional isoform changes in genes whose overall expression was stable (**Figure S2F, Table S1D**). Switching was dominated by alternative transcript start and end sites, with exon skipping the most frequent internal event and intron retention comparatively rare (**Figure 3D**). Top hits included critical proteins involved in innate immune sensing such as IL18BP, USP18, and IRF7 (**Figure 3E**). Isoform switching was predominantly contrast-specific with 66% of switching genes detected in a single contrast, and only six genes contained switching isoforms in all three contrasts (*IREB2, ISG20, NAMPT, SHISA5, TAPBP*, and *VPS26C*). This suggests that FLUAV infection induces extensive host transcriptome isoform remodeling that is largely distinct from the canonical IFN-β response, consistent with virus-driven rewiring of host RNA-processing pathways beyond a small shared antiviral core (**Figure 3F**). Gene ontology analysis of DTU genes identified terms consistent with DTE-derived pathways, including innate immune sensing, viral infection, and defective RIPK1-mediated necroptosis (**Figure 3G**).

**Figure 3.**
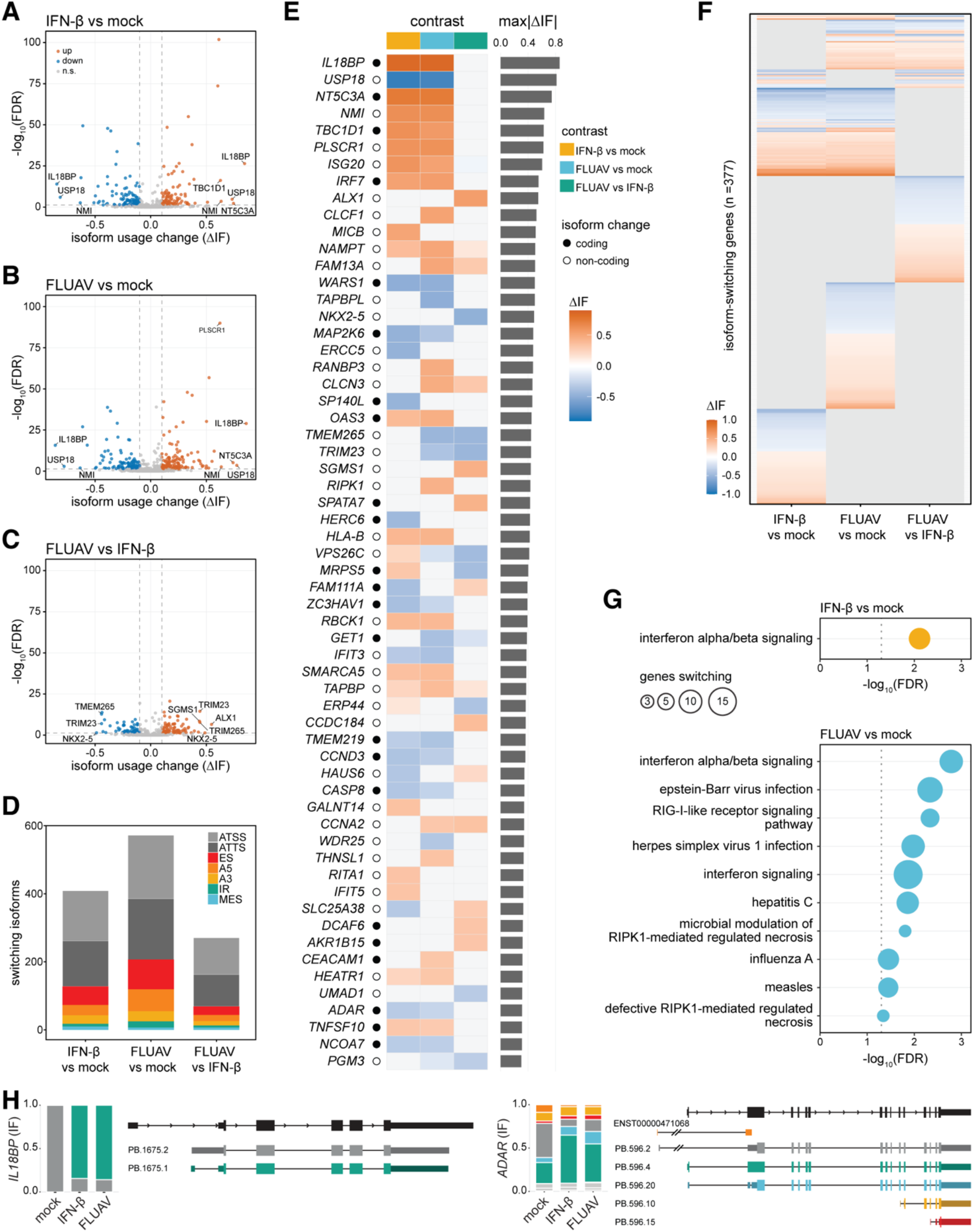
Differential transcript usage analysis reveals genes and pathways that toggle transcript isoforms. Differential transcript usage measures proportionally whether an isoform fraction changes between conditions. **A**-**C**, change in isoform fraction (ΔIF) for transcripts across conditions. Each dot represents a transcript. **D**, Architecture of isoform switching events: alternative transcription start or termination site (ATSS, ATTS), exon skipping (ES), Alternative 5’ or 3’ splice site (A5, A3), intron retention (IR), or multiple exon skipping (MES). **E**-**F**, Top isoform switching genes, ranking the 60 with the highest accumulated ΔIF across contrasts (**E**) or the entire set clustered by treatment (**F**). **G**, Pathways enriched in each of the three contrasts based on gene significance from differential transcript usage. Note that no pathways were enriched for FLUAV vs IFN-β. **H**, Isoform fraction and transcript maps for example genes *IL18BP* and *ADAR*.

We next investigated the impact of these isoform switches on the encoded proteins. Protein-altering switches were concentrated among the largest usage shifts: genes with a coding-level consequence skewed toward high ΔIF values, while the low-magnitude tail was enriched for regulatory (start/end-site) switches. For example, IL18BP binds to IL-18 to limit its pro-inflammatory signaling (56). In mock cells, an intron is retained in the 5’-UTR of IL18BP (PB.1675.2). This intron contains an upstream ORF (uORF) that likely limits IL18BP expression at the basal level (**Figure 3H**). However, IFN-β stimulation or FLUAV infection triggers dramatic removal of this intron, leading to a transcript (PB.1675.1) with a shorter 5’-UTR lacking the uORF, which is expected to result in increased IL18BP expression, possibly to limit the duration of the IL18-driven response (**Figure 3H**). We also identified the well-known isoform switch in ADAR driven by IFN signaling (57). The IFN-insensitive transcript encoding the p110 isoform (PB.596.2) is the most abundant in mock cells. Upon FLUAV infection or IFN-β stimulation, the transcript encoding p150 (PB.596.4), the IFN-induced isoform, became the most abundant, confirming that our experimental approach captures biologically relevant isoform switches during FLUAV infection and innate immune sensing (**Figure 3H**). In conclusion, DTU analysis identified both known and novel isoform switches during FLUAV infection and innate immune sensing.

### Event-level eCLIP integration implicates RNA-binding proteins in immune remodeling and nominates DDX3X at stimulus-specific isoform switches

To interrogate which RBPs control these isoform changes upon FLUAV infection and IFN-β treatment, we turned to ENCODE, which contains rich eCLIP datasets of 232 experiments covering 170 unique RNA-binding proteins across two cell lines (22). Gene-body enrichment analysis of both the full DTU switch gene set and the internal splicing subset yielded no significant RBP associations after Benjamini–Hochberg correction in any of the three contrasts (0 of 232 RBPs, FDR < 0.05), indicating that DTU switch genes are not preferentially targeted by any of the 232 RBPs at the gene-body level (**Figure 4A, S3A**). In contrast, event-level localization analysis identified 16 of 232 RBPs with significant eCLIP peak concentration at internal switch event windows (FDR < 0.05) in one or more contrasts (**Figure 4A, S3A**). During IFN-β signaling, the only RBP identified was DDX3X, a helicase widely involved in innate immune sensing (**Figure 4B**). During FLUAV infection, five RBPs were identified: DDX3X, PRPF8, NCBP2, TIAL1, and U2AF2 (**Figure 4C**). The identification of DDX3X in both contexts confirmed the validity of our experimental approach, as both conditions drive sustained interferon signaling (**Figure S2D)** and would be expected to share RBPs controlling the IFN-β transcriptome. Lastly, comparison of FLUAV infection to IFN-β signaling revealed 13 RBPs differentially engaged during FLUAV infection relative to IFN-β stimulation: AQR, BUD13, DDX3X, DROSHA, IGF2BP2, NCBP2, PABPN1, PPIG, PRPF4, RBFOX2, SF3B4, UCHL5, and ZNF622 (**Figure 4D**). These FLUAV-specific RBPs which show enriched binding at the splicing events are known to have established roles in immune regulation such as myeloid cell recruitment (IGF2BP2; (58)), miRNA-mediated immune control (DROSHA; (59)), and PD-L1 expression-associated splicing programs (RBFOX2; (60)). Thus, the increased engagement of these immune-regulatory RBPs at FLUAV-associated switching events supports a model in which the virus remodels host immune regulation through transcriptome isoform switching, potentially enhancing viral fitness. Similarly, UCHL5 binding was increased in FLUAV relative to IFN-β (**Figure S3B**) and has been implicated in promoting Wnt/β-catenin signaling, a pathway frequently associated with immune suppression (61). This observation further supports a role for virus-induced RBP networks in orchestrating host immune remodeling. Overall, these 16 RBPs encompass spliceosomal proteins (AQR, BUD13, PPIG, PRPF4, SF3B4), proteins involved in ribosome and miRNA biogenesis (DROSHA, ZNF622), and RBPs involved in all steps of the mRNA life cycle (DDX3X, IGF2BP2, NCBP2, PABPN1, RBFOX2) (55).

**Figure 4.**
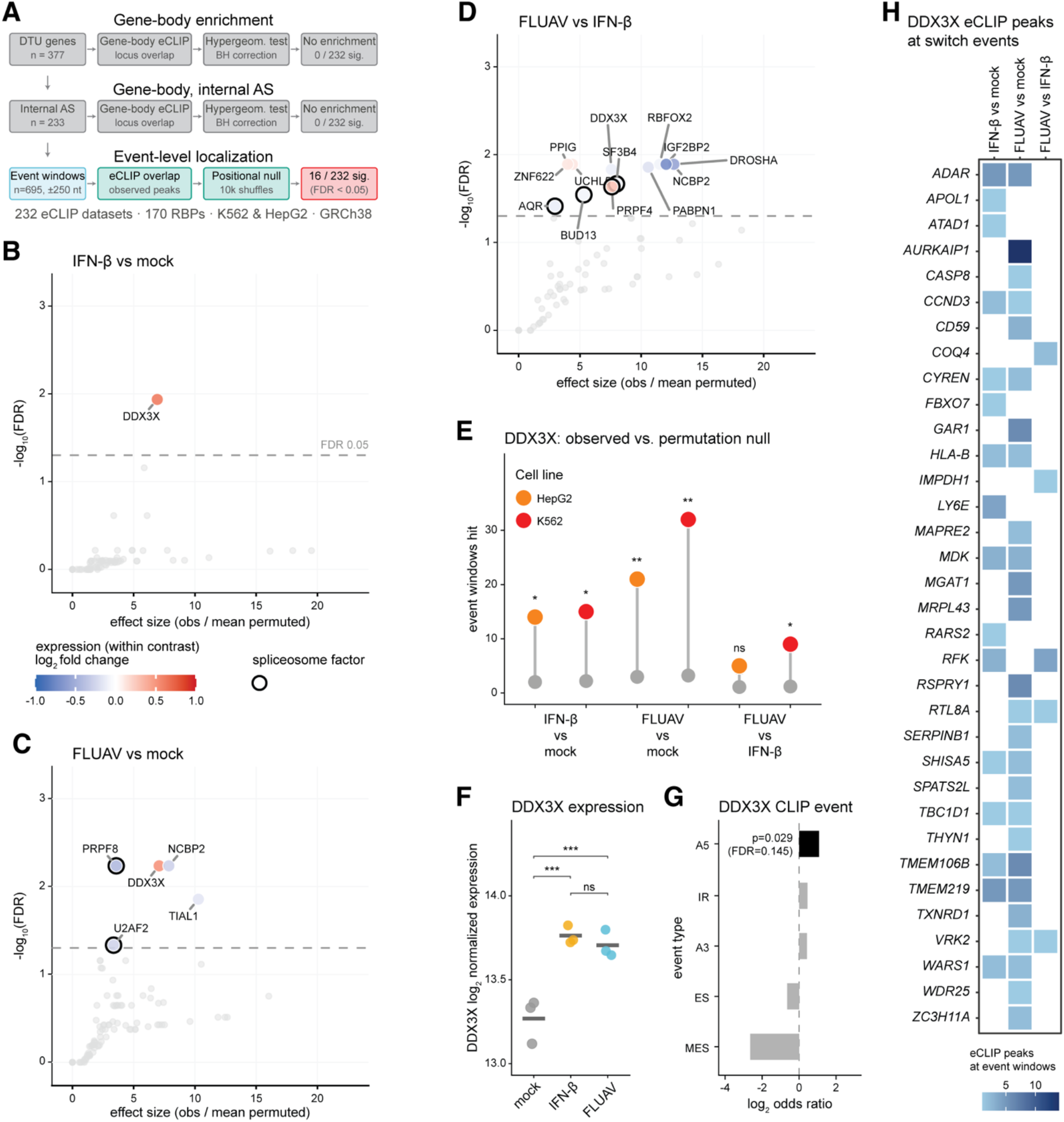
*In silico* analysis of RNA binding proteins targeting genes with differential transcript usage. Enrichment analysis utilized ENCODE enhanced cross-linking immunoprecipitation (eCLIP) datasets of 170 RNA binding proteins (RBPs) from human K562 and HepG2 cell lines. **A**, Three approaches used to identify RBPs with enriched targeting of genes with differential transcript usage (DTU). **B**-**D**, Enrichment for RBPs binding transcripts from DTU genes within splicing event windows (± 250 nucleotides) compared to binding DTU transcripts at randomly shuffled locations (10,000 permutations). Differential expression of significant (FDR < 0.05) RBPs indicated by shading, and spliceosomal factors indicated with thick borders. eCLIP datasets from HepG2 and K562, when both available for the same RBP, were merged. **E**, DDX3X binding within DTU event windows compared to background showing enrichment across cell lines. One-sided empirical P values (+1 pseudocount) were Benjamini-Hochberg corrected across RBPs within each contrast. *, FDR < 0.05; **, FDR < 0.01; ns, not significant. **F**, DDX3X gene-level expression (log2 size-factor-normalized counts summed across SQANTI3 isoforms), points represent biological replicates and the bar marks the group mean. Differences were tested by one-way ANOVA with Tukey HSD post-hoc comparisons. ***, *P* < 0.001; ns, not significant. **G**, DDX3X eCLIP peak preference across DTU switch-event windows by event type (A5, alternative 5′ splice site; A3, alternative 3′ splice site; IR, intron retention; ES, exon skipping; MES, multiple exon skipping), shown as the log_2_ odds ratio of DDX3X-bound versus background (all) event windows; the dashed line at 0 marks no preference (OR = 1). Enrichment was tested per event type by one-sided Fisher’s exact test on deduplicated event windows (n = 340) with Benjamini–Hochberg correction across the five event types, with the exact *P* and FDR annotated for the enriched type (A5). **H**, DDX3X peak counts at DTU switch event windows shown per switch gene (rows) across three contrasts (columns). Fill intensity summed across HepG2 and K562 eCLIP experiments.

DDX3X achieved significance in all three contrasts across both cell lines, suggesting it might play an important role in shaping the transcriptome during innate immune sensing (**Figure 4E**). Moreover, DDX3X expression was significantly increased during IFN-β treatment and FLUAV infection relative to mock, with no additional virus-specific induction beyond the impact of IFN signaling, suggesting it might be an ISG itself regulated at the transcriptional level (**Figure 4F**). DDX3X eCLIP peaks showed nominal enrichment at alternative 5’ splice-site event windows (**Figure 4G**). Peak count analysis revealed that DDX3X bound event windows in 34 unique switch genes across contrasts, with contrast-specific targets including LY6E and RFK upon IFN-β stimulation, consistent with contrast-specific isoform switching rather than constitutive binding (**Figure 4H**).

This pattern extended to all 16 significant RBPs, indicating that binding to regulated targets is stimulus-specific (**Figure S3B**). It is tempting to speculate that the DDX3X eCLIP peaks recovered on the 5’ end of ADAR transcripts control the switch between p110 and p150 (**Figure S3C**). In conclusion, combining DTU analysis with publicly available eCLIP datasets proved invaluable for identifying candidate RBPs associated with transcriptome remodeling during FLUAV infection and IFN-β stimulation, including DDX3X.

## DISCUSSION

In this study, we demonstrated the clear advantage of hybrid sequencing for accurately quantifying complete transcript isoforms and deciphering the shared and distinct transcriptomic changes regulating the cellular response to FLUAV infection or IFN-β stimulation. Using SQANTI3, we robustly profiled isoform changes triggered upon infection or the interferon response and identified genes undergoing isoform switching under these conditions. This dataset is a rich resource for understanding isoform diversity arising upon viral infection or innate immune sensing. Many of the genes that switch isoforms during infection show no significant change in overall expression. Pathway analysis from DTU-defined events highlighted changes in RIPK1 regulation between IFN and FLUAV. Functional RIPK1-mediated necroptosis is a critical anti-viral defense mechanism, suggesting that viral manipulation of RIPK1 transcript splicing (**Fig 3E, Table S1C**) and RIPK1 pathways (**Figure 3G**) can circumvent this host restriction. The biology captured here is therefore distinct from what standard RNA-seq experiments reveal.

Ashraf *et al*. and Thompson *et al*. mapped influenza-induced alternative splicing from short-read data at the level of individual junctions (11,12), and Ueda *et al*. profiled the interferon response at isoform resolution (62). Our study builds on this work in three ways. We compare virus infection directly against interferon stimulation, which separates virus-specific isoform changes from the shared interferon program. Similar to Ueda *et al*., we build the analysis on a hybrid long- and short-read SQANTI3-curated transcriptome, which resolves full-length isoforms and removes reconstruction artifacts before quantification. Finally, we integrate event-level eCLIP to nominate the RBPs positioned at switch windows, such as DDX3X, adding a candidate regulatory layer that expression- or junction-level analyses do not provide. When we intersected the set of validated alternative splicing events from the Ashraf *et al*. and Thompson *at al*. studies with our DTU results, approximately half were not testable because the SQANTI3 long-read transcriptome did not detect them as multi-isoform genes (**Table S1E**) (11,12). This limited testability partly reflects our long-read design. Genes expressed at low level, and those whose minor isoforms fall below Iso-Seq detection, are therefore not represented and do not register as multi-isoform switch candidates. Among the 25 testable genes, 10 reached our effect-size threshold (absolute ΔIF ≥ 0.1) and three cleared both the significance and effect-size gates (NCOA7, IFIT1, and HECTD1). RIPK2 and IRF1 fell just short of significance (FDR = 0.069 and 0.065), while SCD passed the significance gate decisively but not the effect-size threshold. These results highlight that event-level splicing analysis and isoform-proportion analysis measure complementary but distinct aspects of transcriptome remodeling.

Using the ENCODE eCLIP datasets, we identified with high confidence 16 significant RBPs across our three contrasts that represent valuable candidates for understanding isoform regulation during FLUAV infection and IFN stimulation. These significant RBPs can be further classified by stimulus-responsiveness (NCBP2 appears in two groups, up-regulated relative to mock but down-regulated in FLUAV relative to IFN-β): six were stimulus-induced (DDX3X, DROSHA, IGF2BP2, RBFOX2, ZNF622, NCBP2), three were stimulus-repressed (NCBP2, PABPN1, TIAL1), six were constitutive spliceosomal factors (PRPF8, PRPF4, SF3B4, AQR, U2AF2, BUD13), and two showed no differential expression (PPIG, UCHL5). The identification of spliceosomal proteins, and notably PRPF8, as critical factors controlling isoform diversity during FLUAV infection was somewhat expected and strongly validates our approach. PRPF8 is the core protein of the U5 snRNP, one of the three snRNPs of the spliceosome critical for catalyzing the splicing reaction (63,64). PRPF8 was previously identified in siRNA screens as a critical factor for FLUAV replication (65,66), and FLUAV infection was further shown to increase PRPF8 levels to bolster viral replication (67). PRPF8 was also linked to innate immune sensing alongside the snRNP proteins EFTUD2 and SNRNP200 (52,68–72). RBPs for FLUAV-associated isoform-switching events converged on host immune regulation. This suggests that viral-induced isoform remodeling is not restricted to generic RNA-processing programs but may engage post-transcriptional networks capable of reshaping host immune states. However, our identification of RBPs is limited to publicly available eCLIP datasets. As the number of publicly available datasets grows, additional integration and identification of relevant RBPs during viral infection and innate immune sensing will become possible.

Among the RBPs we identified, DDX3X, an X-linked helicase with widespread roles in the mRNA life cycle (including transcription, splicing, translation, stability, and stress granule assembly), played a central role in regulating isoform diversity during FLUAV infection and innate immune sensing (73). DDX3X plays an important role in innate immune sensing, though some results are contradictory. DDX3X is phosphorylated by TBK1 and recruited to the IFN-β promoter for efficient type-I interferon induction (74,75). In another study, DDX3X was shown to restrict the type I interferon response during mammarenavirus infection and following RIGI agonist stimulation, suggesting a context-dependent function (76). During FLUAV infection, DDX3X is required for NLRP3 inflammasome activation (77). In the absence of NS1, an important FLUAV immune evasion protein, DDX3X also enhanced the assembly of stress granules and type I interferon responses (77). The involvement of DDX3X in NLRP3 inflammasome activation was also independently validated by another group (78).

Among the targets shown to be directly bound by DDX3X (**Figure S3B**) are isoforms with well-known functions in the innate immune response. One such target is ADAR p150, an interferon-inducible A-to-I RNA editing enzyme that is crucial for preventing autoinflammation and limiting innate immune signaling, notably during FLUAV infection (57,79,80). This suggests DDX3X might act as a broad negative regulator of innate immune responses at the post-transcriptional level. Discrepancies across studies may arise from cellular context, including the presence or absence of its Y-linked homolog DDX3Y, which may also shape DDX3X activity (73). We observed a modest increase in mRNA abundance of both DDX3X and DDX3Y upon FLUAV infection and IFN-β stimulation, suggesting this increase is driven by the cellular antiviral state (**Figure 4F**). However, this study is the first to link DDX3X to isoform diversity under these conditions. Additional work should aim to determine whether DDX3X exerts its effect through splicing or stability, whether it also plays roles in alternative transcription start sites and polyadenylation, and how these isoform changes contribute to the cellular antiviral state. Together, these results raise the possibility that DDX3X acts as a post-transcriptional rheostat of innate immune activation. DDX3X is an interferon-induced RNA helicase that may limit the amplitude or duration of the immune response by promoting isoform switching at key negative regulators like ADAR. Testing this model constitutes a natural next step for experimental follow-up.

A potential concern is that the ENCODE eCLIP datasets were generated from unstimulated K562 and HepG2 cells rather than from virus-infected or cytokine-stimulated A549 cells. Two features of our results argue that this homeostatic binding is still informative. First, binding is target- and stimulus-specific: DDX3X and the other 15 significant RBPs concentrate at the switch-event windows of the genes that switch under each stimulus (**Figure 4H, S3B**), so pre-existing peaks mark the very transcripts that later undergo stimulus-specific remodeling. Second, the significant RBP hits are concordant across two independent cell lines, which suggests that these binding relationships are conserved across cellular contexts and are likely relevant in A549 cells. The 170 RBPs covered by ENCODE eCLIP represent approximately 11% of the full human RBP census of 1,542 (55). The identification of 16 significant RBPs from this partial sampling underscores how much of the regulatory landscape remains to be explored, and expanding eCLIP coverage will likely reveal additional candidates controlling isoform diversity during viral infection and innate immune sensing.

In conclusion, we leveraged hybrid sequencing using SQANTI3 to generate the first hybrid long- and short-read transcript map of A549 cells upon FLUAV infection and IFN-β stimulation, allowing us to decipher the shared and distinct transcriptomic changes upon cellular perturbation. Taking advantage of the ENCODE datasets and our high-resolution transcript map, we identified RBPs associated with these isoform changes and suggest DDX3X as a candidate regulator of the transcriptome during FLUAV infection and IFN-β stimulation. This dataset and the methodology developed here constitute a rich resource for understanding isoform diversity arising upon viral infection, innate immune sensing, or other cellular perturbations.

## Acknowledgements

This work was supported by the American Lung Association (ERPALA2023). We want to thank Dr. Martin Bisaillon (Université de Sherbrooke), Dr. Kristen Lynch (University of Pennsylvania), and Dr. Tommy Alain (Children’s Hospital of Eastern Ontario Research Institute, University of Ottawa) for their scientific input and logistical support.

## Author contributions

Conceptualization: GS, SFB

Methodology: SB, RHB, AG, GS, SFB

Formal Analysis: SB, RHB, AG, SFB

Investigation: SB, RHB, AG, SFB

Writing – Original Draft: SB & SFB

Writing – Review & Editing: SB, GS, SFB

Visualization: SFB

Funding Acquisition: SFB

Supervision: GS, SFB

## METHODS

### Hybrid sequencing

Long-read (PacBio, Iso-Seq) and short-read (Illumina, 150 bp paired-end) RNA sequencing data were generated from A549 cells under three conditions: mock, interferon-beta treatment (IFN-β; 250 U/ml, 8 h), and influenza A virus infection (FLUAV; A/WSN/33 (H1N1), MOI 0.02, 24 h). Library preparation and sequencing were described previously (21,50). Short-read sequencing comprised nine libraries (three conditions, three biological replicates each; BioProject PRJNA667475). For Iso-Seq, three condition-specific primers were used during library prep and the biological replicates within each condition were pooled and sequenced across two SMRT cells (BioProject PRJNA1423310).

Subreads from each SMRT cell were processed into circular consensus sequence (CCS) reads with ccs v6.4.0 (`--min-rq 0.9`). Full-length reads were identified and demultiplexed by primer pair with lima v2.9.0 in Iso-Seq mode (`--isoseq --peek-guess`), which orients reads 5′ to 3′ and removes barcodes. Poly(A) tails and artificial concatemers were removed with isoseq refine v4.0.0 (`--require-polya`) to yield full-length non-chimeric (FLNC) reads. FLNC reads from all conditions and both SMRT cells were merged and clustered with isoseq cluster2 v4.0.0 (`--use-qvs`). Clustered transcripts were aligned to the reference genome with pbmm2 v1.13.1 (`-- preset ISOSEQ --sort`) and collapsed into unique isoforms with isoseq collapse (`--do-not-collapse-extra-5exons`). Per-sample FLNC counts were retained by supplying the FLNC file list to the collapse step.

Collapsed isoforms were polished with SQANTI3 v5.2.1 (40). The reference used for both long-read mapping and SQANTI3 was a combined human and FLUAV assembly and annotation, referred to as humanplusflu (GRCh38 and A/WSN/33 (H1N1)) (81). In the quality-control step, each isoform was classified by its splice-junction match to the reference annotation and scored on a panel of structural quality descriptors.

Orthogonal evidence was supplied for this step: transcription start sites were assessed against refTSS v3.1 (82), 3′ ends against a human poly(A) motif list, per-isoform full-length support from the Iso-Seq FLNC counts, and splice-junction support and coverage from the nine short-read RNA-seq libraries, which were provided as FASTQ and aligned internally with STAR. Potential artifacts were removed with the SQANTI3 rules filter using a custom rules set. Isoforms were discarded for intra-priming (high genomic adenine content immediately downstream of the transcript termination site), reverse-transcriptase template switching, non-canonical splice junctions lacking short-read support, and related quality failures. To avoid discarding genuine transcripts that fail orthogonal validation, the SQANTI3 rescue module was applied in automatic mode. The reference annotation was first quality-controlled with the same data and criteria; full splice match artifacts were then replaced by their corresponding reference transcripts, and discarded novel isoforms were reassigned to the best-supported reference or long-read target. Rescue recovered 853 transcripts. The filtered and rescued models constitute the final curated human and influenza transcriptome used for short-read quantification.

### Differential transcript expression (DTE) analysis

Short-read RNA-seq was processed with the nf-core/rnaseq pipeline v3.14.0 (83) under Nextflow v24.04.3. Reads were adapter- and quality-trimmed with fastp v0.23.4, aligned to the genome with STAR v2.7.9a (84), and quantified at the transcript level with Salmon v1.10.1 (85); per-sample quantifications were summarized into a transcript count matrix with tximport (86). The nine libraries were quantified against two references using identical pipeline parameters. The default reference combined humanplusflu with Ensembl release 112 annotation. The SQANTI3 reference combined humanplusflu and the rescued transcriptome described above. DTE was assessed for each reference with DESeq2 v1.44.0 (Bioconductor 3.19) with R v4.4.1 (51). For each transcript-level count matrix, counts were modeled with the design `∼ condition`, and Wald tests were performed for three contrasts: IFN-β versus mock, FLUAV versus mock, and FLUAV versus IFN-β. Transcripts were called significantly differentially expressed (sig UP or DN) at a Benjamini-Hochberg adjusted *P*-value < 0.01 and an absolute log_2_ fold change > 0.58 (a 1.5-fold change). Nominal transcripts (nom UP or DN) failed multiple-testing correction: *P* < 0.01 and |log_2_ fold change| > 0.58, but FDR ≥ 0.01. For the reported counts, transcripts not expressed in the conditions being compared (median count below five) were excluded. Transcript identifiers were annotated with gene symbols using biomaRt against Ensembl GRCh38.p14 (87). Oppositely regulated transcripts (ORTs) were identified from sig UP/DN transcripts. For each multi-isoform gene, the numbers of significantly up- and down-regulated transcripts were tallied per contrast, and a gene was called oppositely regulated in a contrast when it contained at least one significantly up-regulated and at least one significantly down-regulated transcript in that same contrast.

### RT-qPCR

RT-PCR wAS performed as described before (52,88–90). The complete list of primers used in this study is available in **Table S1F**. Briefly, reverse transcription was done in a 20 μL reaction using 1 μg of RNA, random primers (S1330, NEB), and in-house purified MMLV reverse transcriptase (Plateforme de purification des protéines, Université de Sherbrooke). cDNA was diluted to 5 ng/μL following the RT step in water. Quantitative PCR (qPCR) reactions were prepared in 10 μl in 96-well plates with 5 μL of in-house prepared 2X SYBR qPCR Master Mix (Plateforme de purification des protéines, Université de Sherbrooke), 2 μL of cDNA (10 ng), and 0.4 μM of each primer. qPCR reactions were performed on a C1000 Touch Thermal Cycler (Bio-Rad) and data acquired with a CFX96 Dx Real-Time PCR Detection Systems for In Vitro Diagnostics (Bio-Rad). Analysis was done with CFX-Maestro 2.0 (Bio-Rad). PUM1 was used as the housekeeping gene for normalization. For all qPCR experiments, control reactions (no RT) were performed in the absence of cDNA for each primer pair and confirmed to be negative.

### Pathway enrichment analysis

Significant transcripts from DTE analysis moving in both directions per contrast were retained throughout. For each contrast, significant genes were submitted to g:Profiler against KEGG and Reactome, with up- and downregulated sets submitted separately so that opposing regulation within a pathway would not cancel. The union of significant terms across all six directional queries defined the pathway universe (216 pathways). Each pathway was expanded to its complete member-gene complement (Reactome via reactome.db, KEGG via KEGGREST) rather than only the differentially expressed genes that defined it, so that the downstream test evaluates whether a pathway as a whole shifts. Every measured gene was collapsed to its highest-baseMean transcript, carrying all three contrasts’ log_2_ fold changes, and the full measured gene set was retained as the competitive background (measured genes, 7,976; pathway members, 4,166; background, 3,810). For each contrast, every pathway’s measured member genes were compared against all other measured genes with a competitive Wilcoxon ranksum test, Benjamini-Hochberg corrected across pathways (significance threshold 0.05; pathways with fewer than three measured members were not tested). The test was run twice to detect coordinated directional regulation (signed log_2_ fold change) and magnitude of perturbation regardless of direction (absolute log_2_ fold change). For reporting and visualization we removed pathways with >300 measured member genes and collapsed pathways that had nested members as determined by Jaccard ≥ 0.7 (for example overlapping Reactome subtrees).

RNA-binding proteins (RBPs) enriched in DTE analysis were defined in (55) (1,542 genes) with the following exclusions: ribosome-associated genes (RNA target = “ribosome”), histones (removed by gene name), and proteasome subunits. Retained RBPs were ranked by membership in the 216-term pathway universe derived from over-representation analysis of differential transcript expression (DTE; KEGG and Reactome). Three ranking metrics were computed: the raw number of pathways containing each RBP; an inverse-term-size weighted score (Σ 1/term size) that down-weights large, generic pathways; and the weighted score restricted to non-mega pathways (term size ≤ 300), used as the primary ranking. Gene-level expression was computed by summing isoform counts per gene and applying a blind variance-stabilizing transformation (DESeq2 vst), shown per biological replicate across mock, IFN-β, and FLUAV. Only RBPs detected in the sq3 transcriptome (above a summed-count filter) were retained in the figure.

Pathways overrepresented from differential transcript usage (DTU) analysis (below) were defined as follows. A foreground gene list of 377 isoform switchers was compared to a background gene list within the DTU-testable universe (4,786 genes with >=2 isoforms passing the DTU pre-filter). Switch genes were tested for enrichment of KEGG and Reactome pathways (GO:BP reported separately) with g:Profiler (gprofiler2) with Benjamini-Hochberg FDR correction at 0.05. FDR was used in place of g:Profiler’s native g:SCS, which is calibrated for the whole-genome domain and is over-conservative against a small custom background. Switching was assessed direction-agnostically at the gene level, so each contrast contributed a single foreground.

### Differential transcript usage (DTU) analysis

Differential transcript usage was assessed with IsoformSwitchAnalyzeR (91). Salmon quantifications (three biological replicates per condition) against the SQANTI3 transcriptome were imported with inferential replicates, and a switchAnalyzeRlist was constructed for three contrasts. Low-expressed isoforms and single-isoform genes were removed (preFilter), and differential isoform fraction (ΔIF) was tested per contrast using the DEXSeq backbone. Genes were considered significant isoform switchers when the gene-level switch qvalue (a Benjamini-Hochberg adjusted *P*-value) < 0.05 and at least one isoform showed an absolute ΔIF ≥ 0.1 in any contrast (377 genes). For annotation, termed consequence, ORFs were assigned from SQANTI3 GTF CDS records; protein domains were annotated with pfam_scan/HMMER against Pfam-A, coding potential with CPAT (human model, coding cutoff 0.725), and local splicing events and switch consequences with analyzeAlternativeSplicing and analyzeSwitchConsequences. Consequence directions oriented to the mock vs stimulus comparison.

### ENCODE eCLIP data analysis

eCLIP peak BED files for 232 experiments covering 170 unique RNA-binding proteins (RBPs; K562 and HepG2 cell lines, GRCh38) were obtained from the ENCODE Project portal. Three independent intersection tests were performed against DTU switch events identified by IsoformSwitchAnalyzeR (q < 0.05, |ΔIF| ≥ 0.1). For gene-body enrichment of all switch genes (n = 377), eCLIP peaks were intersected with gene locus coordinates using bedtools intersect; a one-sided hypergeometric test was applied per RBP against 4,409 non-switching expressed background genes, with Benjamini-Hochberg correction across all 232 RBPs per contrast. A second test was performed restricting the foreground to genes with internal-splicing (exon-skipping, alternative splice sites, or intron retention; n = 233). Event-level intersection selected windows from internal-splicing genes (695 windows from 233 genes; windows ±250 nt flanking each regulated splicing event). A positional permutation null was constructed by shuffling the gene-body-filtered eCLIP peaks 10,000 times within switch gene body coordinates using bedtools shuffle; the empirical one-sided *P*-value was computed with a +1/+1 pseudocount (92) and Benjamini-Hochberg-corrected across RBPs per contrast. RBPs with fewer than 20 gene-body peaks were excluded. DDX3X event-type preference was assessed by one-sided Fisher’s exact test on geometrically deduplicated event windows (n = 340) with Benjamini-Hochberg correction across five event types. Gene-level expression of RBPs was derived from SQANTI3 pipeline DESeq2 size-factor-normalized counts (log_2_-transformed) by summing isoform counts per gene per sample. To quantify RBP binding at individual switch genes, eCLIP peaks for each significant RBP were intersected with per-contrast internal event windows using bedtools intersect -c, counting peaks per window. Counts were summed across cell lines per (RBP, gene, contrast) to produce gene-level peak counts.

### Data and code availability

Raw sequencing data are available under BioProjects PRJNA1423310 (Iso-Seq) and PRJNA667475 (shortread; accessions SRR12775100 to SRR12775108). The curated transcriptome annotation, the transcript count matrices, and all analysis code (Iso-Seq processing, SQANTI3 curation, the nf-core/rnaseq configuration, and the DESeq2 differential expression workflow) are available at https://github.com/sfbaker/boudreault-sqanti-fluavifn and archived at Zenodo (DOI: 10.5281/zenodo.22830056).

## SUPPLEMENTAL FIGURES

**Figure S1.**
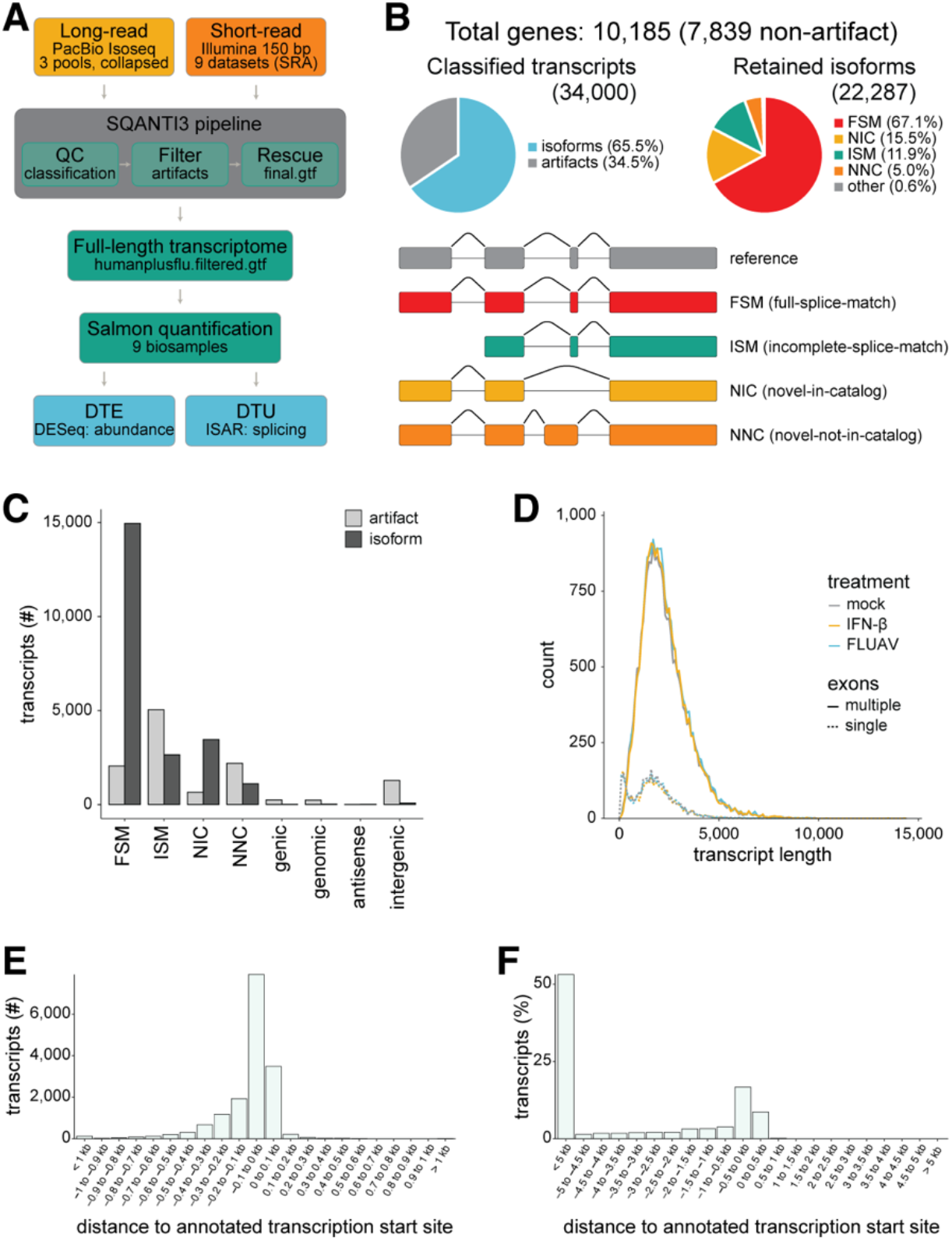
Analysis of genome-wide full-length transcript regulation. **A**, Analysis pipeline overview. Biological triplicate RNA samples from A549 cells (mock, IFN treated or infected with influenza A virus) were used for library prep and long- or shortread RNA-sequencing. Long-read RNA-sequencing (Isoseq, PacBio) results, polished with short-read data (SQANTI3) informed the identity of full-length transcripts present in these experimental conditions. Short-read data can then be aligned using salmon and data analyzed by abundance (DESeq) or transcript use and splicing (ISAR). **B**, SQANTI3 analysis after polishing removed transcript artifacts (24.5%), and annotated remaining isoforms as variants of reference transcript models. **C**, Number of transcripts by type and whether removed as artifact by SQANTI3 or annotated as isoforms. **D**, Distribution of transcript length for single or multi exon genes across treatment conditions. **E**-**F**, Distribution of transcription start sites of transcripts from full-splice-match (**E**) vs incomplete-splice-match (**F**) categories relative to annotated start sites.

**Figure S2.**
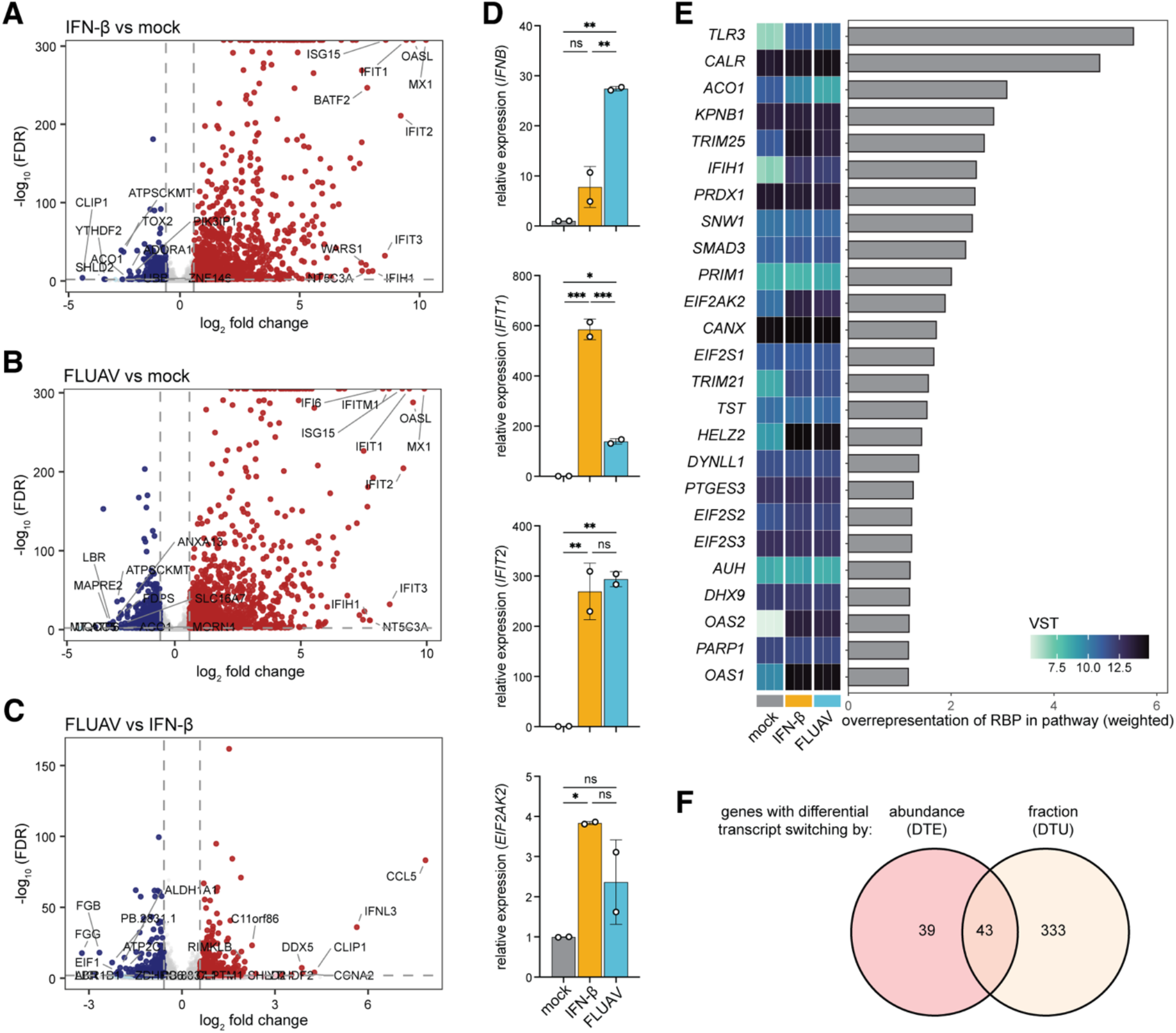
Differential transcript expression shows similar gene and RNA binding protein enrichment between influenza infection and interferon treatment. **A**-**C**, Differentially expressed transcripts between indicated conditions. Red, upregulated; blue, downregulated. All transcripts plotted, but only one representative circle annotated per gene indicating the top 10 most differentially expressed. **D**, Relative expression of target genes per condition measured by RT-qPCR. **E**, Genes encoding RNA binding proteins enriched from pathway analysis in **Fig. 2 A-C** ranked by weighted overrepresentation in pathways. Heatmap shows gene variance-stabilized counts (VST) with biological replicates per treatment condition. **F**, Overlap of genes with differentially expressed isoforms identified by differential transcript expression (DTE; DESeq2, custom script) or differential transcript usage (DTU; IsoformSwitchAnalyzeR).

**Figure S3.**
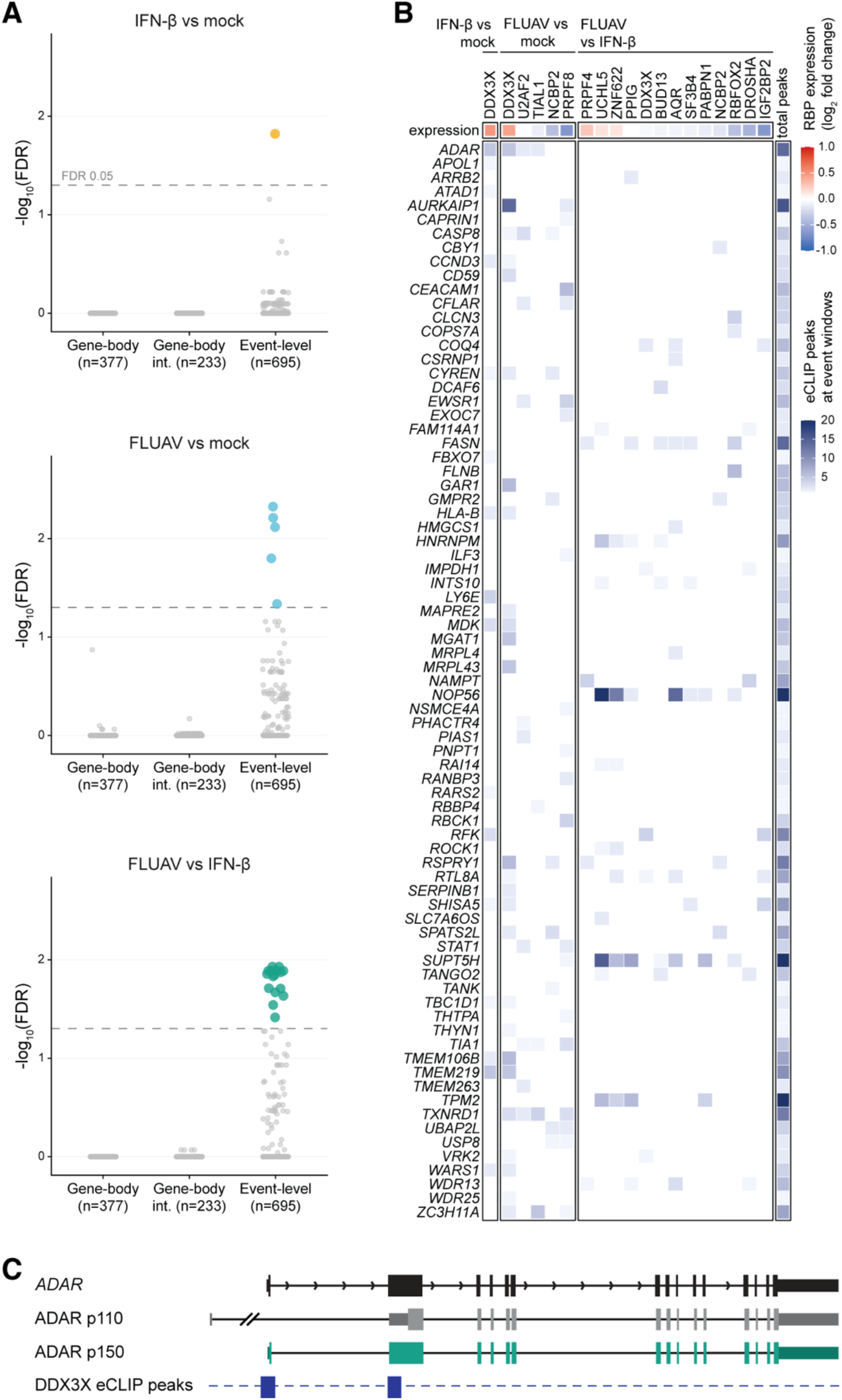
RNA binding proteins with enriched enhanced cross-linking and immunoprecipitation (eCLIP) peaks. **A**, Full RNA binding protein (RBP) analysis of ENCODE data using three intersection approaches. Event-level analysis discovered RBPs with eCLIP peak enrichment. **B**, RBP eCLIP peaks (columns) within event-window-containing genes (rows). **C**, DDX3X eCLIP peaks at the exons controlling *ADAR* p110 versus p150 expression.

## REFERENCES

1. Fensterl V, Chattopadhyay S, Sen GC. No Love Lost Between Viruses and Interferons. Annu Rev Virol (2015) 2:549–572. doi: 10.1146/annurev-virology-100114-055249

2. Fensterl V, Sen GC. Interferon-Induced Ifit Proteins: Their Role in Viral Pathogenesis. J Virol (2015) 89:2462–2468. doi: 10.1128/JVI.02744-14

3. Sen GC, Sarkar SN. The interferon-stimulated genes: targets of direct signaling by interferons, double-stranded RNA, and viruses. Curr Top Microbiol Immunol (2007) 316:233–250.

4. Teijaro JR. Type I interferons in viral control and immune regulation. Current Opinion in Virology (2016) 16:31–40. doi: 10.1016/j.coviro.2016.01.001

5. Rutkowski AJ, Erhard F, L’Hernault A, Bonfert T, Schilhabel M, Crump C, Rosenstiel P, Efstathiou S, Zimmer R, Friedel CC, et al. Widespread disruption of host transcription termination in HSV-1 infection. Nature Communications (2015) 6:7126. doi: 10.1038/ncomms8126

6. Jia X, Yuan S, Wang Y, Fu Y, Ge Y, Ge Y, Lan X, Feng Y, Qiu F, Li P, et al. The role of alternative polyadenylation in the antiviral innate immune response. Nat Commun (2017) 8:14605. doi: 10.1038/ncomms14605

7. Boudreault S, Roy P, Lemay G, Bisaillon M. Viral modulation of cellular RNA alternative splicing: A new key player in virus–host interactions? Wiley Interdiscip Rev RNA (2019) 10:e1543. doi: 10.1002/wrna.1543

8. Mann JT, Riley BA, Baker SF. All differential on the splicing front: Host alternative splicing alters the landscape of virus-host conflict. Seminars in Cell & Developmental Biology (2023) 146:40–56. doi: 10.1016/j.semcdb.2023.01.013

9. Begg BE, Ferretti MB, Tracey MA, Lynch KW. Viral Modulation of Host Splicing. Annual Review of Virology (2025) 12:203–222. doi: 10.1146/annurev-virology-092623-102539

10. Boudreault S, Martenon-Brodeur C, Caron M, Garant J-M, Tremblay M-P, Armero VES, Durand M, Lapointe E, Thibault P, Tremblay-Létourneau M, et al. Global Profiling of the Cellular Alternative RNA Splicing Landscape during Virus-Host Interactions. PLoS One (2016) 11:e0161914. doi: 10.1371/journal.pone.0161914

11. Ashraf U, Benoit-Pilven C, Navratil V, Ligneau C, Fournier G, Munier S, Sismeiro O, Coppée J-Y, Lacroix V, Naffakh N. Influenza virus infection induces widespread alterations of host cell splicing. NAR Genomics and Bioinformatics (2020) 2: doi: 10.1093/nargab/lqaa095

12. Thompson MG, Dittmar M, Mallory MJ, Bhat P, Ferretti MB, Fontoura BM, Cherry S, Lynch KW. Viral-induced alternative splicing of host genes promotes influenza replication. eLife (2020) 9:e55500. doi: 10.7554/eLife.55500

13. Thompson MG, Muñoz-Moreno R, Bhat P, Roytenberg R, Lindberg J, Gazzara MR, Mallory MJ, Zhang K, García-Sastre A, Fontoura BMA, et al. Co-regulatory activity of hnRNP K and NS1-BP in influenza and human mRNA splicing. Nature Communications (2018) 9:2407. doi: 10.1038/s41467-018-04779-4

14. Nemeroff ME, Barabino SML, Li Y, Keller W, Krug RM. Influenza Virus NS1 Protein Interacts with the Cellular 30 kDa Subunit of CPSF and Inhibits 3′ End Formation of Cellular Pre-mRNAs. Molecular Cell (1998) 1:991–1000. doi: 10.1016/S1097-2765(00)80099-4

15. Hale BG, Steel J, Medina RA, Manicassamy B, Ye J, Hickman D, Hai R, Schmolke M, Lowen AC, Perez DR, et al. Inefficient Control of Host Gene Expression by the 2009 Pandemic H1N1 Influenza A Virus NS1 Protein. Journal of Virology (2010) 84:6909–6922. doi: 10.1128/jvi.00081-10

16. Steidle S, Martínez-Sobrido L, Mordstein M, Lienenklaus S, García-Sastre A, Stäheli P, Kochs G. Glycine 184 in Nonstructural Protein NS1 Determines the Virulence of Influenza A Virus Strain PR8 without Affecting the Host Interferon Response. Journal of Virology (2010) 84:12761–12770. doi: 10.1128/jvi.00701-10

17. Gaucherand L, Porter BK, Levene RE, Price EL, Schmaling SK, Rycroft CH, Kevorkian Y, McCormick C, Khaperskyy DA, Gaglia MM. The Influenza A Virus Endoribonuclease PA-X Usurps Host mRNA Processing Machinery to Limit Host Gene Expression. Cell Reports (2019) 27:776–792.e7. doi: 10.1016/j.celrep.2019.03.063

18. Gaucherand L, Iyer A, Gilabert I, Rycroft CH, Gaglia MM. Cut site preference allows influenza A virus PA-X to discriminate between host and viral mRNAs. Nat Microbiol (2023) 8:1304–1317. doi: 10.1038/s41564-023-01409-8

19. Khaperskyy DA, Schmaling S, Larkins-Ford J, McCormick C, Gaglia MM. Selective Degradation of Host RNA Polymerase II Transcripts by Influenza A Virus PA-X Host Shutoff Protein. PLOS Pathogens (2016) 12:e1005427. doi: 10.1371/journal.ppat.1005427

20. Bauer DLV, Tellier M, Martínez-Alonso M, Nojima T, Proudfoot NJ, Murphy S, Fodor E. Influenza Virus Mounts a Two-Pronged Attack on Host RNA Polymerase II Transcription. Cell Reports (2018) 23:2119–2129.e3. doi: 10.1016/j.celrep.2018.04.047

21. Baker SF, Meistermann H, Tzouros M, Baker A, Golling S, Polster JS, Ledwith MP, Gitter A, Augustin A, Javanbakht H, et al. Alternative splicing liberates a cryptic cytoplasmic isoform of mitochondrial MECR that antagonizes influenza virus. PLOS Biology (2022) 20:e3001934. doi: 10.1371/journal.pbio.3001934

22. Van Nostrand EL, Freese P, Pratt GA, Wang X, Wei X, Xiao R, Blue SM, Chen J-Y, Cody NAL, Dominguez D, et al. A large-scale binding and functional map of human RNA-binding proteins. Nature (2020) 583:711–719. doi: 10.1038/s41586-020-2077-3

23. Panthi A, Lynch KW. RNA processing in innate immunity: regulation by RNA-binding proteins. Trends in Biochemical Sciences (2025) 50:610–621. doi: 10.1016/j.tibs.2025.04.004

24. Boudreault S, Rivera-Lopez Y, Ferretti MB, Tracey MA, Bonner J, Jacobs BL, Lynch KW. Nonsense-mediated decay controls a negative feedback loop in innate immune sensing. Proceedings of the National Academy of Sciences (2026) 123:e2517478123. doi: 10.1073/pnas.2517478123

25. Zhao W, Liu M, Kirkwood KL. p38α Stabilizes Interleukin-6 mRNA via Multiple AU-richElements*. Journal of Biological Chemistry (2008) 283:1778–1785. doi: 10.1074/jbc.M707573200

26. Naora H, Young IG. Mechanisms Regulating the mRNA Levels of Interleukin-5 and Two Other Coordinately Expressed Lymphokines in the Murine T Lymphoma EL4.23. Blood (1994) 83:3620–3628. doi: 10.1182/blood.V83.12.3620.3620

27. Braun RM, Ferretti MB, Lee JS, Miller J, Castellana L, Acuña J, Whig K, Dohnalová L, Descamps HC, Huber AS, et al. Selective decay of interferon mRNAs controls antiviral immunity and innate memory. Molecular Cell (2026) 86:3565–3584.e10. doi: 10.1016/j.molcel.2026.07.022

28. Wagner AR, Scott HM, West KO, Vail KJ, Fitzsimons TC, Coleman AK, Carter KE, Watson RO, Patrick KL. Global Transcriptomics Uncovers Distinct Contributions From Splicing Regulatory Proteins to the Macrophage Innate Immune Response. Front Immunol (2021) 12: doi: 10.3389/fimmu.2021.656885

29. Scott HM, Smith MH, Coleman AK, Armijo KS, Chapman MJ, Apostalo SL, Wagner AR, Watson RO, Patrick KL. Serine/arginine-rich splicing factor 7 promotes the type I interferon response by activating Irf7 transcription. Cell Reports (2024) 43: doi: 10.1016/j.celrep.2024.113816

30. West KO, Scott HM, Torres-Odio S, West AP, Patrick KL, Watson RO. The Splicing Factor hnRNP M Is a Critical Regulator of Innate Immune Gene Expression in Macrophages. Cell Reports (2019) 29:1594–1609.e5. doi: 10.1016/j.celrep.2019.09.078

31. Zhou B, Wu F, Han J, Qi F, Ni T, Qian F. Exploitation of nuclear protein SFPQ by the encephalomyocarditis virus to facilitate its replication. Biochemical and Biophysical Research Communications (2019) 510:65–71. doi: 10.1016/j.bbrc.2019.01.032

32. Pozzi B, Bragado L, Mammi P, Torti MF, Gaioli N, Gebhard LG, García Solá ME, Vaz-Drago R, Iglesias NG, García CC, et al. Dengue virus targets RBM10 deregulating host cell splicing and innate immune response. Nucleic Acids Res (2020) 48:6824–6838. doi: 10.1093/nar/gkaa340

33. Lee N, Pimienta G, Steitz JA. AUF1/hnRNP D is a novel protein partner of the EBER1 noncoding RNA of Epstein-Barr virus. RNA (2012) 18:2073–2082. doi: 10.1261/rna.034900.112

34. Batra R, Stark TJ, Clark E, Belzile J-P, Wheeler EC, Yee BA, Huang H, Gelboin-Burkhart C, Huelga SC, Aigner S, et al. RNA-binding protein CPEB1 remodels host and viral RNA landscapes. Nat Struct Mol Biol (2016) 23:1101. doi: 10.1038/nsmb.3310

35. Kneller ELP, Connor JH, Lyles DS. hnRNPs Relocalize to the Cytoplasm following Infection with Vesicular Stomatitis Virus. J Virol (2009) 83:770–780. doi: 10.1128/JVI.01279-08

36. Brunetti JE, Scolaro LA, Castilla V. The heterogeneous nuclear ribonucleoprotein K (hnRNP K) is a host factor required for dengue virus and Junín virus multiplication. Virus Research (2015) 203:84–91. doi: 10.1016/j.virusres.2015.04.001

37. Barnhart MD, Moon SL, Emch AW, Wilusz CJ, Wilusz J. Changes in cellular mRNA stability, splicing, and polyadenylation through HuR protein sequestration by a cytoplasmic RNA virus. Cell Rep (2013) 5:909–917. doi: 10.1016/j.celrep.2013.10.012

38. Bentley DR, Balasubramanian S, Swerdlow HP, Smith GP, Milton J, Brown CG, Hall KP, Evers DJ, Barnes CL, Bignell HR, et al. Accurate whole human genome sequencing using reversible terminator chemistry. Nature (2008) 456:53–59. doi: 10.1038/nature07517

39. Lu H, Giordano F, Ning Z. Oxford Nanopore MinION Sequencing and Genome Assembly. genom proteom bioinform (2016) 14:265–279. doi: 10.1016/j.gpb.2016.05.004

40. Pardo-Palacios FJ, Arzalluz-Luque A, Kondratova L, Salguero P, Mestre-Tomás J, Amorín R, Estevan-Morió E, Liu T, Nanni A, McIntyre L, et al. SQANTI3: curation of long-read transcriptomes for accurate identification of known and novel isoforms. Nat Methods (2024) 21:793–797. doi: 10.1038/s41592-024-02229-2

41. Dai H, Deng P, Wang K, Lv B, Yang Z, Kong C, Xie Z, Jia J, Xia C Du F, et al. Alternative splicing is associated with tissue differentiation, subgenome divergence, and agronomic trait regulation in hexaploid wheat. Plant Physiol (2026) 201:kiag211. doi: 10.1093/plphys/kiag211

42. Tang K-W, Alaei-Mahabadi B, Samuelsson T, Lindh M, Larsson E. The landscape of viral expression and host gene fusion and adaptation in human cancer. Nature Communications (2013) 4:2513. doi: 10.1038/ncomms3513

43. Jeon S, Kukreja B, Jang J, Liang Y, Hahm A, Turcke CE, Yim YS, Kalish BT. Maternal immune activation perturbs the brain epitranscriptome. Brain, Behavior, and Immunity (2026) 137:106804. doi: 10.1016/j.bbi.2026.106804

44. Bresnahan ST, Yong HEJ, Nemani A, Wu WH, Lopez S, Chan JKY, White F, Jacques P-É, Hivert M-F, Chan S-Y, et al. Long-read assembly reveals vast transcriptional complexity in the placenta associated with metabolic and endocrine function. Nat Commun (2026) 17:4795. doi: 10.1038/s41467-026-71303-4

45. Wheeler EC, Van Nostrand EL, Yeo GW. Advances and challenges in the detection of transcriptome-wide protein-RNA interactions. Wiley Interdiscip Rev RNA (2018) 9: doi: 10.1002/wrna.1436

46. Van Nostrand EL, Huelga SC, Yeo GW. Experimental and Computational Considerations in the Study of RNA-Binding Protein-RNA Interactions. Adv Exp Med Biol (2016) 907:1–28. doi: 10.1007/978-3-319-29073-7_1

47. Nussbacher JK, Batra R, Lagier-Tourenne C, Yeo GW. RNA-binding proteins in neurodegeneration: Seq and you shall receive. Trends Neurosci (2015) 38:226–236. doi: 10.1016/j.tins.2015.02.003

48. Larson RS ed. “Ontology-Driven Approaches to Analyzing Data in Functional Genomics - Springer.,” Methods in Molecular Biology. Humana Press (2006) http://link.springer.com.ezproxy.usherbrooke.ca/protocol/10.1385%2F1-59259-964-8%3A67 [Accessed April 22, 2015]

49. Thomas PD, Mi H, Lewis S. Ontology annotation: mapping genomic regions to biological function. Current Opinion in Chemical Biology (2007) 11:4–11. doi: 10.1016/j.cbpa.2006.11.039

50. Nath H, Arndt A, Mann JT, Baker SF. Tandem split-GFP influenza A viruses for sensitive and accurate replication analyses. Microbiol Spectr (2025) 14:e02772-25. doi: 10.1128/spectrum.02772-25

51. Love MI, Huber W, Anders S. Moderated estimation of fold change and dispersion for RNA-seq data with DESeq2. Genome Biol (2014) 15:550. doi: 10.1186/s13059-014-0550-8

52. Boudreault S, Lemay G, Bisaillon M. U5 snRNP Core Proteins Are Key Components of the Defense Response against Viral Infection through Their Roles in Programmed Cell Death and Interferon Induction. Viruses (2022) 14:2710. doi: 10.3390/v14122710

53. Su G, Chen Y, Li X, Shao J-W. Virus versus host: influenza A virus circumvents the immune responses. Front Microbiol (2024) 15: doi: 10.3389/fmicb.2024.1394510

54. Song M, Wang Y, Zhang H, Ji X, Sun S, Fang Z, Ying J, Li Q, Chen J. RNA splicing in cancer cell death regulation: shedding light on the molecular mechanisms and potential clinical applications. Cell Death Dis (2026) doi: 10.1038/s41419-026-08687-0

55. Gerstberger S, Hafner M, Tuschl T. A census of human RNA-binding proteins. Nature Reviews Genetics (2014) 15:829. doi: 10.1038/nrg3813

56. Kim S-H, Eisenstein M, Reznikov L, Fantuzzi G, Novick D, Rubinstein M, Dinarello CA. Structural requirements of six naturally occurring isoforms of the IL-18 binding protein to inhibit IL-18. Proceedings of the National Academy of Sciences (2000) 97:1190–1195. doi: 10.1073/pnas.97.3.1190

57. Hu S-B, Heraud-Farlow J, Sun T, Liang Z, Goradia A, Taylor S, Walkley CR, Li JB. ADAR1p150 prevents MDA5 and PKR activation via distinct mechanisms to avert fatal autoinflammation. Molecular Cell (2023) 83:3869–3884.e7. doi: 10.1016/j.molcel.2023.09.018

58. Li J, Jiang Y, Ma M, Wang L, Jing M, Yang Z, Wang L, Qiu Q, Song R, Pu Y, et al. IGF2BP2 Shapes the Tumor Microenvironment by Regulating Monocyte and Macrophage Recruitment in Bladder Cancer. Cancer Medicine (2024) 13:e70506. doi: 10.1002/cam4.70506

59. Shapiro JS, Schmid S, Aguado LC, Sabin LR, Yasunaga A, Shim JV, Sachs D, Cherry S, tenOever BR. Drosha as an interferon-independent antiviral factor. Proceedings of the National Academy of Sciences (2014) 111:7108–7113. doi: 10.1073/pnas.1319635111

60. Cho N, Kim S-Y, Lee S-G, Park C, Choi S, Kim E-M, Kim KK. Alternative splicing of PBRM1 mediates resistance to PD-1 blockade therapy in renal cancer. EMBO J (2024) 43:7. doi: 10.1038/s44318-024-00262-7

61. Bao L, Wu Y, Ren Z, Huang Y, Jiang Y, Li K, Xu X, Ye Y, Gui Z. Comprehensive pan-cancer analysis indicates UCHL5 as a novel cancer biomarker and promotes cervical cancer progression through the Wnt signaling pathway. Biol Direct (2024) 19:139. doi: 10.1186/s13062-024-00588-6

62. Ueda MT, Inamo J, Miya F, Shimada M, Yamaguchi K, Kochi Y. Functional and dynamic profiling of transcript isoforms reveals essential roles of alternative splicing in interferon response. Cell Genom (2024) 4:100654. doi: 10.1016/j.xgen.2024.100654

63. Newman AJ. The role of U5 snRNP in pre-mRNA splicing. EMBO J (1997) 16:5797–5800. doi: 10.1093/emboj/16.19.5797

64. Wood KA, Eadsforth MA, Newman WG, O’Keefe RT. The Role of the U5 snRNP in Genetic Disorders and Cancer. Front Genet (2021) 12:636620.

65. Brass AL, Huang I-C, Benita Y, John SP, Krishnan MN, Feeley EM, Ryan BJ, Weyer JL, Weyden L van der, Fikrig E, et al. The IFITM Proteins Mediate Cellular Resistance to Influenza A H1N1 Virus, West Nile Virus, and Dengue Virus. Cell (2009) 139:1243–1254. doi: 10.1016/j.cell.2009.12.017

66. Karlas A, Machuy N, Shin Y, Pleissner K-P, Artarini A, Heuer D, Becker D, Khalil H, Ogilvie LA, Hess S, et al. Genome-wide RNAi screen identifies human host factors crucial for influenza virus replication. Nature (2010) 463:818–822. doi: 10.1038/nature08760

67. Yang C-H, Li H-C, Shiu Y-L, Ku T-S, Wang C-W, Tu Y-S, Chen H-L, Wu C-H, Lo S-Y. Influenza A virus upregulates PRPF8 gene expression to increase virus production. Arch Virol (2017) 162:1223–1235. doi: 10.1007/s00705-016-3210-3

68. Tremblay N, Baril M, Chatel-Chaix L, Es-Saad S, Park AY, Koenekoop RK, Lamarre D. Spliceosome SNRNP200 Promotes Viral RNA Sensing and IRF3 Activation of Antiviral Response. PLoS Pathog (2016) 12:e1005772. doi: 10.1371/journal.ppat.1005772

69. De Arras L, Laws R, Leach SM, Pontis K, Freedman JH, Schwartz DA, Alper S. Comparative Genomics RNAi Screen Identifies Eftud2 as a Novel Regulator of Innate Immunity. Genetics (2014) 197:485–496. doi: 10.1534/genetics.113.160499

70. Hu P, Li Y, Zhang W, Liu R, Peng L, Xu R, Cai J, Yuan H, Feng T, Tian A, et al. The Spliceosome Factor EFTUD2 Promotes IFN Anti-HBV Effect through mRNA Splicing. Mediators of Inflammation (2023) 2023:e2546278. doi: 10.1155/2023/2546278

71. Zhu C, Xiao F, Hong J, Wang K, Liu X, Cai D, Fusco DN, Zhao L, Jeong SW, Brisac C, et al. EFTUD2 Is a Novel Innate Immune Regulator Restricting Hepatitis C Virus Infection through the RIG-I/MDA5 Pathway. J Virol (2015) 89:6608–6618. doi: 10.1128/JVI.00364-15

72. Zhu C, Xiao F, Lin W. EFTUD2 on innate immunity. Oncotarget (2015) 6:32313–32314.

73. Owens MC, Yanas A, Liu KF. Sex chromosome-encoded protein homologs: current progress and open questions. Nat Struct Mol Biol (2024) 31:1156–1166. doi: 10.1038/s41594-024-01362-y

74. Soulat D, Bürckstümmer T, Westermayer S, Goncalves A, Bauch A, Stefanovic A, Hantschel O, Bennett KL, Decker T, Superti-Furga G. The DEAD-box helicase DDX3X is a critical component of the TANK-binding kinase 1-dependent innate immune response. EMBO J (2008) 27:2135–2146. doi: 10.1038/emboj.2008.126

75. Schloer S, Hennesen J, Rueschpler L, Zamzamy M, Flomm F, Ip WH, Pirosu A, Dobner T, Altfeld M. The host cell factor DDX3 mediates sex dimorphism in the IFNα response of plasmacytoid dendritic cells upon TLR activation. Pharmacological Research (2025) 216:107764. doi: 10.1016/j.phrs.2025.107764

76. Chiale C, Thelen F, Zuniga E. DDX3X suppresses RIG-I dependent type I Interferon production and promotes Mammarenavirus growth. J Immunol (2024) 212:0133_5959. doi: 10.4049/jimmunol.212.supp.0133.5959

77. Kesavardhana S, Samir P, Zheng M, Malireddi RKS, Karki R, Sharma BR, Place DE, Briard B, Vogel P, Kanneganti T-D. DDX3X coordinates host defense against influenza virus by activating the NLRP3 inflammasome and type I interferon response. J Biol Chem (2021) 296:100579. doi: 10.1016/j.jbc.2021.100579

78. Kienes I, Bauer S, Gottschild C, Mirza N, Pfannstiel J, Schröder M, Kufer TA. DDX3X Links NLRP11 to the Regulation of Type I Interferon Responses and NLRP3 Inflammasome Activation. Front Immunol (2021) 12: doi: 10.3389/fimmu.2021.653883

79. Vogel OA, Han J, Liang C-Y, Manicassamy S, Perez JT, Manicassamy B. The p150 Isoform of ADAR1 Blocks Sustained RLR signaling and Apoptosis during Influenza Virus Infection. PLOS Pathogens (2020) 16:e1008842. doi: 10.1371/journal.ppat.1008842

80. Jiao H, Wachsmuth L, Wolf S, Lohmann J, Nagata M, Kaya GG, Oikonomou N, Kondylis V, Rogg M, Diebold M, et al. ADAR1 averts fatal type I interferon induction by ZBP1. Nature (2022) 607:776–783. doi: 10.1038/s41586-022-04878-9

81. Russell AB, Trapnell C, Bloom JD. Extreme heterogeneity of influenza virus infection in single cells. eLife (2018) 7:e32303. doi: 10.7554/eLife.32303

82. Abugessaisa I, Noguchi S, Hasegawa A, Kondo A, Kawaji H, Carninci P, Kasukawa T. refTSS: A Reference Data Set for Human and Mouse Transcription Start Sites. Journal of Molecular Biology (2019) 431:2407–2422. doi: 10.1016/j.jmb.2019.04.045

83. Ewels PA, Peltzer A, Fillinger S, Patel H, Alneberg J, Wilm A, Garcia MU, Di Tommaso P, Nahnsen S. The nf-core framework for community-curated bioinformatics pipelines. Nat Biotechnol (2020) 38:276–278. doi: 10.1038/s41587-020-0439-x

84. Dobin A, Davis CA, Schlesinger F, Drenkow J, Zaleski C, Jha S, Batut P, Chaisson M, Gingeras TR. STAR: ultrafast universal RNA-seq aligner. Bioinformatics (2013) 29:15–21. doi: 10.1093/bioinformatics/bts635

85. Patro R, Duggal G, Love MI, Irizarry RA, Kingsford C. Salmon provides fast and bias-aware quantification of transcript expression. Nat Methods (2017) 14:417–419. doi: 10.1038/nmeth.4197

86. Soneson C, Love MI, Robinson MD. Differential analyses for RNA-seq: transcript-level estimates improve gene-level inferences. (2016) doi: 10.12688/f1000research.7563.2

87. Durinck S, Spellman PT, Birney E, Huber W. Mapping identifiers for the integration of genomic datasets with the R/Bioconductor package biomaRt. Nat Protoc (2009) 4:1184–1191. doi: 10.1038/nprot.2009.97

88. Boudreault S, Armero VES, Scott MS, Perreault J-P, Bisaillon M. The Epstein-Barr virus EBNA1 protein modulates the alternative splicing of cellular genes. Virol J (2019) 16:29. doi: 10.1186/s12985-019-1137-5

89. Boudreault S, Durand M, Martineau C-A, Perreault J-P, Lemay G, Bisaillon M. Reovirus μ2 protein modulates host cell alternative splicing by reducing protein levels of U5 snRNP core components. Nucleic Acids Res (2022) 50:5263–5281. doi: 10.1093/nar/gkac272

90. Boudreault S, Martineau C-A, Faucher-Giguère L, Abou-Elela S, Lemay G, Bisaillon M. Reovirus μ2 Protein Impairs Translation to Reduce U5 snRNP Protein Levels. International Journal of Molecular Sciences (2023) 24:727. doi: 10.3390/ijms24010727

91. Han C, Gilis J, Delgado EI, Clement L, Vitting-Seerup K. IsoformSwitchAnalyzeR v2: analysis of functional isoform changes in long-read and single-cell sequencing data. NAR Genom Bioinform (2026) 8:lqag098. doi: 10.1093/nargab/lqag098

92. Phipson B, Smyth GK. Permutation P-values Should Never Be Zero: Calculating Exact P-values When Permutations Are Randomly Drawn. Statistical Applications in Genetics and Molecular Biology (2010) 9: doi: 10.2202/1544-6115.1585

